# Tumor-Associated EDA-FN-Enriched Matrix Instructs Macrophage Behavior

**DOI:** 10.64898/2026.05.14.725237

**Authors:** Ghazal Bashiri, Elizabeth Bakare, Jessica Longstreth, Marshall S. Padilla, Karin Wang

## Abstract

**Introduction:** Cancer progression is driven not only by tumor cells but also by interactions between the extracellular matrix (ECM), stromal cells, and immune cells within the tumor microenvironment (TME). Cancer-associated fibroblasts (CAFs) are major drivers of ECM remodeling, assembling ECM with aberrant organization. Extra domain A fibronectin (EDA-FN), a cellular FN containing an extra type III domain, is upregulated in the TME. EDA-FN regulates cellular behavior and has been associated with poor patient prognosis. Macrophages are among the most abundant immune cells within the TME, where they contribute to TME remodeling and inflammation to promote cancer cell invasion and metastasis. However, how tumor-associated matrix-specific cues regulate macrophage behavior remains largely understudied.

**Purpose:** Here, we developed a fibroblast-derived matrix platform that captures the structural imprint of tumor-associated EDA-enriched matrices and investigated how matrix-specific cues regulate macrophage behavior in the absence of ongoing soluble factor cues.

**Method:** Human mammary fibroblasts (HMFs) preconditioned in incubated low-serum media (lNC, or control) and MDA-MB231 metastatic breast cancer cell-conditioned media (mTCM) were cultured on polyacrylamide gels of 2 kPa and 20 kPa, respectively, followed by decellularization. Matrix organization, including fiber alignment, width, and intrafibrillar spacing, was quantified from confocal images. Decellularized EDA-FN-enriched matrices were subsequently reseeded with macrophages to assess macrophage morphology, phenotype, and matrix interactions.

**Results:** The combined effects of tumor-derived soluble factors and pathological stiffness induced a CAF-like phenotype in HMFs, accompanied by cytoskeletal reorganization and microarchitectural alterations of EDA-FN-enriched matrices. Tumor-associated matrices exhibited increased alignment, narrower fiber width, and enlarged intrafibrillar spacing compared to control matrices. These aberrant, tumor-associated matrix-derived features were associated with altered macrophage behavior, including heterogeneous morphology, enhanced localized EDA-FN matrix loss beneath the cell body, and a hybrid phenotype with a shift toward a CD206-dominant profile.

**Conclusions:** These findings demonstrate the feasibility of obtaining EDA-FN-enriched matrices to isolate matrix-specific cues for investigating macrophage-ECM interactions. Furthermore, this platform can be leveraged to identify matrix-targeting therapeutic approaches for modulating macrophage function within the TME.

## Introduction

Cancer-related mortality is largely driven by tumor invasion and metastasis to distant organs [1–5]. Notably, human breast cancer progression is not solely dependent on tumor cells, but is regulated by continuous crosstalk between tumor cells and their surrounding microenvironment [6]. The tumor microenvironment (TME) is a highly complex system comprising multiple cell types, including cancer cells, stromal cells, and immune cells, as well as acellular components, such as extracellular matrix (ECM) and signaling molecules [6–9]. The ECM is a complex three-dimensional network of proteins and glycans that not only provides cells with structural support and mechanical cues, but also serves as a reservoir for sequestration of soluble signaling molecules to regulate various cellular behaviors such as adhesion, differentiation, and migration [10–12].

ECM remodeling alters the deposition and organization of ECM components [13]. Although this process is crucial for normal tissue development and homeostasis, under pathological conditions, such as cancer, it becomes dysregulated [4, 13]. Accordingly, dysregulated ECM remodeling generates an abnormal microenvironment by establishing a self-reinforcing cell-matrix interaction feedback loop that promotes cancer progression [14].

Fibronectin (FN) is a multidomain mechanosensitive protein that contains binding sites for cell surface receptors such as integrins and other ECM molecules, thereby regulating various cell functions and contributing to the assembly of other ECM components [15–18]. The peptide sequence arginine–glycine–aspartic acid–serine (RGD) and its adjacent synergy site (PHSRN) are integrin-binding domains located on FNIII10 and FNIII9, respectively, mediating cell adhesion and cytoskeletal organization [19–21]. FN exists in both plasma-soluble and cellular-derived forms [18, 22]. The extra domain A (EDA) FN is an alternatively spliced isoform of cellular FN that contains an additional type III domain [18]. EDA-FN interacts with integrins α9β1, α4β1, α4β7, and α5β1 [23–25], and is highly upregulated in tumor stroma, where it has been associated with poor patient prognosis [12, 26–29].

Cancer-associated fibroblasts (CAFs) represent the most abundant stromal cells within the TME in both primary and metastatic tumors [7, 26, 27]. CAFs play a prominent role in developing tumor progression and metastasis through both biochemical and mechanical signaling [28]. CAFs secrete cytokines and growth factors that promote cancer cell proliferation, angiogenesis, and tumor cell invasion, while recruiting tumor-promoting immune cells to establish an immunosuppressive microenvironment [10, 29, 30]. In addition to biochemical signaling, CAFs are major regulators of ECM deposition, assembly, and remodeling [14, 27]. Notably, CAFs have also been associated with synthesizing and incorporating the cellular isoform of FN containing the EDA domain into the ECM stroma of various tumors [31, 32]. In cancer and tissue fibrosis, CAFs excessively deposit ECM proteins such as FN and collagen, and assemble them into a 3D-structure with aberrant organization [7, 15, 22, 33, 34]. CAF-derived ECM is often characterized by enhanced stiffness, density, and conformational changes of fibrillar structures such as FN and collagen within the TME. Such abnormally remodeled ECM can further release previously sequestered matrix-bound growth factors or disease-specific fragments [35, 36]. In turn, fibroblasts sense these biochemical and mechanical alterations and respond by differentiating into CAFs to maintain a tumor-promoting microenvironment [37].

In tumors, approximately 5-40% of the tumor mass consists of tumor-associated macrophages (TAMs) [38] TAMs display a heterogeneous population with distinct functional phenotypes that exist along a spectrum and dynamically transition in response to biochemical and mechanical cues from their microenvironment, with pro-inflammatory, anti-tumor, and anti-inflammatory, pro-tumor phenotypes representing the two extremes of this spectrum [39–44]. Macrophages interact with the ECM through integrins [23, 45, 46]. Macrophages have been implicated in enhancing ECM deposition and play a prominent role in ECM remodeling through the secretion of matrix metalloproteinases (MMPs), thereby enabling matrix degradation and reorganization [47, 48]. Thus, macrophages significantly contribute to angiogenesis, tumor cell migration, and metastasis [47–51]. Notably, macrophage-matrix interactions are bidirectional. Physical cues from the ECM, including composition and microarchitectural organization, have been shown to regulate macrophage cytoskeletal organization and morphology, to influence their phenotypes and functions, such as migration and ECM remodeling [52–55].

In this context, FN and EDA-FN have been shown to regulate macrophage behavior via different receptor-mediated and signaling pathways [56–58]. For example, FN has been shown to promote macrophage polarization toward an anti-inflammatory phenotype through α5β1 integrin signaling in breast cancer [59], while the RGD domain of FN has been shown to induce TAM-like polarization of monocytes with elevated MMP-9 activity, highlighting the role of macrophage-matrix interactions in ECM remodeling [60]. In addition, the EDA domain of FN has been identified as a damage-associated molecular pattern (DAMP) ligand for innate immune receptors, including Toll-like receptors 4 (TLR4) [24, 61]. And EDA-FN-TLR4 interactions have been implicated in multiple pathological conditions, such as metastatic cancer and chronic fibrosis [24, 58, 62–64]. EDA-FN has also been reported to promote anti-inflammatory responses in macrophages and support tumor progression through binding to α5β1 integrin [65].

Together, these studies highlight the importance of macrophage-matrix interactions in tumor progression. However, how macrophages interact and respond to tumor-associated matrix-specific cues has yet to be explored. Specifically, how the microarchitectural organization of EDA-FN-enriched matrices regulates macrophage function within the tumor microenvironment remains poorly understood. Therefore, this study aimed to develop a tumor-associated human mammary fibroblast (HMF)-derived matrix platform, in which HMFs are exposed to the combined impact of MDA-MB-231 metastatic breast cancer cell-conditioned media and pathological substrate stiffness to generate a decellularized tumor-associated microenvironment (Fig. 1). These cell-derived matrices are subsequently used to investigate how EDA-FN-enriched tumor matrices regulate macrophage behavior in the absence of ongoing soluble factor cues.

**Fig. 1.**
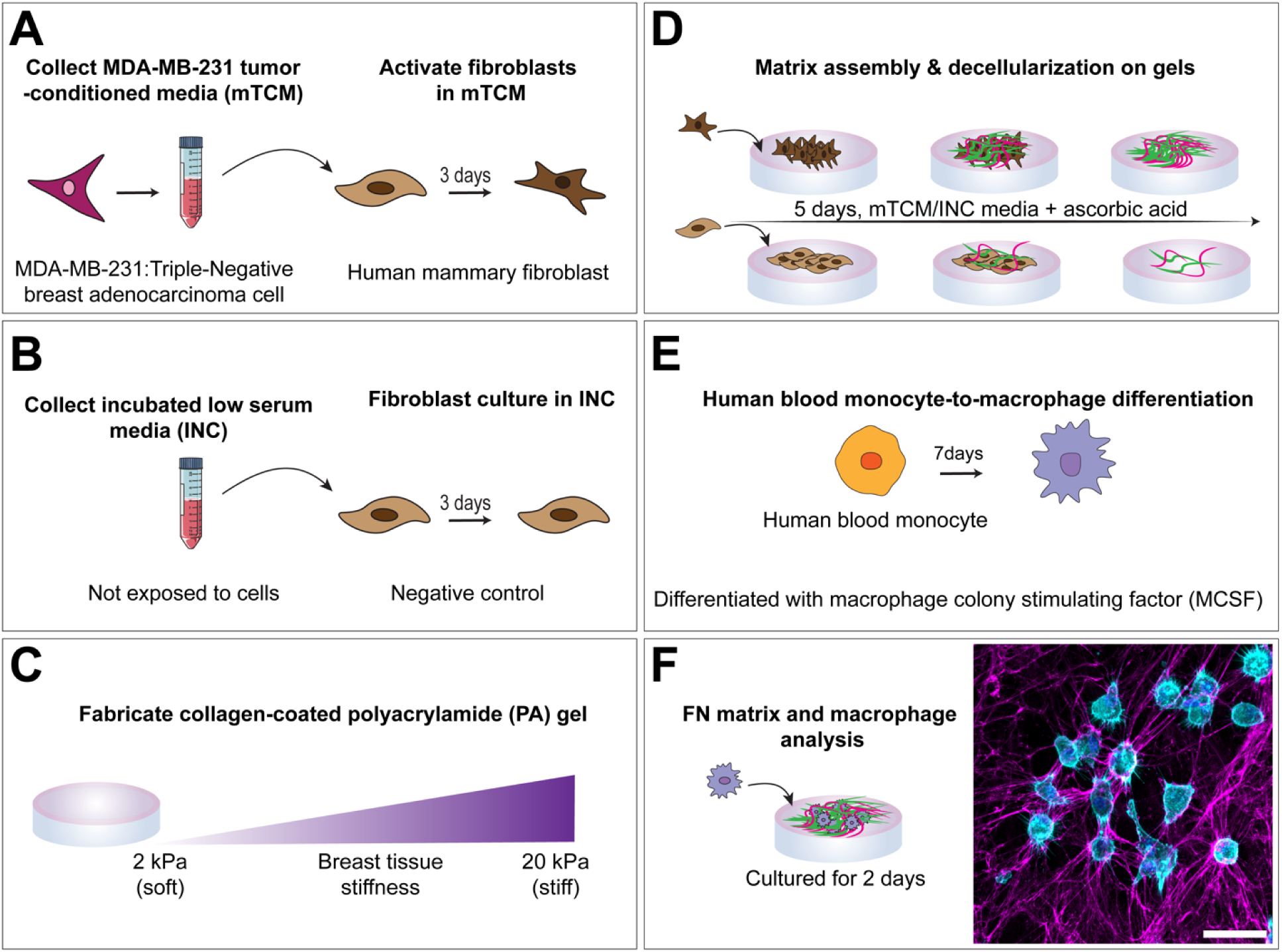
Schematic workflow for developing a platform to study macrophage-EDA-FN-enriched tumor matrix interactions. **A** MDA-MB-231, a highly metastatic triple-negative breast cancer cell line, was used to isolate tumor-conditioned media (mTCM). Human mammary fibroblasts (HMFs) were conditioned in mTCM for 3 days to differentiate into cancer-associated fibroblast (CAF)-like cells. **B** Incubated low-serum media (INC), not exposed to cells, was used to culture HMFs for 3 days as a negative control. **C** Polyacrylamide gels were fabricated with stiffnesses of either ∼2 kPa (soft) or ∼20 kPa (stiff) to recapitulate normal and tumor breast tissue, respectively, and coated with collagen type I. **D** mTCM-conditioned HMFs were seeded on stiff gels, while INC-conditioned HMFs were seeded on soft gels. Each were fed the corresponding conditioned media, mTCM or INC, containing ascorbic acid to promote matrix maturation, every other day for 5 days. On day 5, cells were extracted while the matrices were kept intact. **E** Fresh human blood-derived monocytes were differentiated into macrophages using macrophage colony-stimulating factor (MCSF) over 7 days. **F** Decellularized matrices (magenta) were either fixed and immunostained for imaging and microarchitectural analysis or re-seeded with macrophages (cyan) to study macrophage-EDA-FN-enriched tumor matrix interactions. Scale bar, 10 µm. The figure was made in Adobe Illustrator

## Materials and Methods

### Cell Lines and Culture

Human mammary fibroblasts (HMFs), obtained from ScienCell Research Laboratories (7630), were cultured in Dulbecco’s Modified Eagle Medium (DMEM) supplemented with 10% fetal bovine serum (FBS) and 1% penicillin–streptomycin. Passages 5-6 were used for experiments. MDA-MB-231 cells were obtained from the American Type Culture Collection (ATCC, HTB-26) and cultured in DMEM supplemented with 10% FBS and 1% penicillin–streptomycin. Passages 5-12 were used for experiments. Primary human peripheral blood monocyte from healthy donors, purchased from the University of Pennsylvania Human Immunology Core, were cultured in Roswell Park Memorial Institute (RPMI) 1640 medium supplemented with 10% heat-inactivated human serum (HS) and 1% penicillin–streptomycin.

### MDA-MB-231 Tumor-Conditioned Medium (mTCM) and Incubated Low-Serum Medium (INC) Preparation

Triple-negative MDA-MB-231 breast cancer cells were cultured in high-serum medium (DMEM supplemented with 10% FBS and 1% penicillin–streptomycin). mTCM was collected when cells reached 80-85% confluency. 24 h before mTCM collection, cells were fed low-serum media (DMEM supplemented with 1% FBS and 1% penicillin–streptomycin) and incubated at 37 °C to induce the release of tumor-derived soluble factors rather than promote cell growth. INC was generated by incubating low-serum media at 37 °C for 24 h without any cell exposure and used as the control media. The volume of mTCM and INC collected was normalized to cell number (per 7 million cells). The collected mTCM and INC were concentrated to 10% of their original volume using Amicon^@^ Ultra centrifugal filters (3 kDa MWCO) and centrifuged for 30 min at 4000 rpm at 4 °C. Concentrated mTCM and INC were stored at - 20 °C for future conditioning experiments.

### Enzyme-Linked Immunosorbent Assay (ELISA)

The concentration of human transforming growth factor ß1(TGF-ß1) in mTCM and INC medium was measured by TGF-ß1 DuoSet ELISA kit (R&D Systems, DY240), according to the manufacturer’s instructions. Briefly, clear 96-well plates (ThermoFisher Scientific) were coated overnight with the capture antibody, then washed and blocked. Latent TGF-ß1, in mTCM and INC medium, was activated by acidification followed by neutralization. Captured TGF-ß1 molecules were then detected using a biotinylated detection antibody and streptavidin–HRP, followed by color development. Absorbance was measured at 450 nm with wavelength correction at 540 nm using an Infinite 200 Pro microplate reader (TECAN), and TGF-ß1 concentrations were calculated from a standard curve.

### Polyacrylamide Gel (PA) Fabrication and Collagen Functionalization

#### Coverslip and dish treatment preparation

18-mm glass coverslips were rinsed with ethanol and dried at room temperature. They were then coated with a siliconizing reagent (Sigmacoat; Sigma-Aldrich, SL2-100ML) to make the surface hydrophobic. 35-mm dishes with a 20-mm glass-bottom inset (CellVis, D35-20-1.5-N) were treated with 1N H_2_SO_4_ for 20 min, followed by three washes with deionized water and air drying. The dishes were then treated with 0.1 M NaOH for 20 min, followed by three washes with deionized water and air drying. The dishes were subsequently treated with 3-(Trimethoxysilyl) propyl methacrylate (TMSPMA; Sigma-Aldrich, 440159-100M) for 1 h at room temperature, followed by 3 washes with pure ethanol, three washes with deionized water, and air drying.

#### PA gel fabrication

To model normal and metastatic breast tissue stiffnesses, PA gels with stiffness values of approximately 2 kPa (soft) and 20 kPa (stiff) were fabricated [66–68]. Gel stiffness was controlled by polymerizing acrylamide monomer (40% w/v; Bio-Rad, 1610140) and bis-acrylamide (2% w/v; Bio-Rad, 1610142) in deionized water using ammonium persulfate (APS; Bio-Rad, 1610700) and Tetramethylethylenediamine (TEMED; Bio-Rad, 1610800). The final acrylamide concentration was maintained constant at 8%, while the bis-acrylamide concentration was varied to achieve the desired stiffness. The stiffness of the PA gels was previously characterized using atomic force microscopy (AFM) by Young et al. [67]. To fabricate the gels, 24 µl of each PA solution was pipetted onto each treated glass inset, and Sigmacoat-treated coverslips were gently placed on top to form a gel sandwich. Soft and stiff gels were polymerized for 25 min and 20 min, respectively, in a desiccator. After polymerization, 2 ml of deionized water was added to each gel, and the gels were stored overnight at 4 °C for hydration. Table 1 shows the concentrations and volumes of reagents used for the fabrication of soft and stiff gels.

**Table 1.**
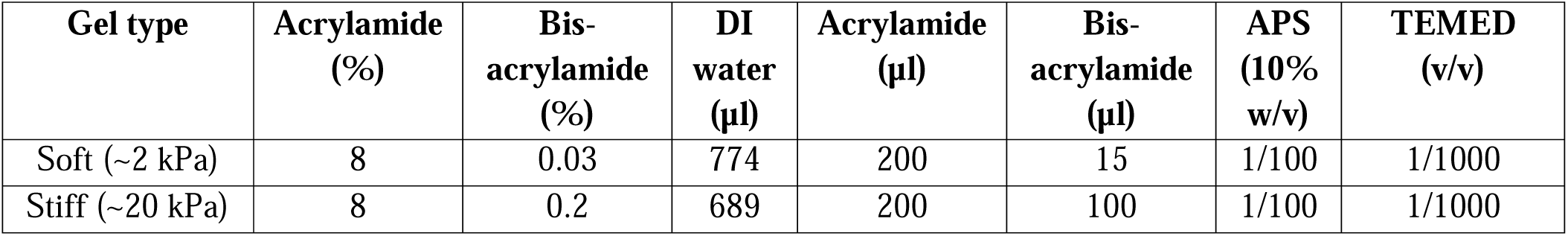
Composition of PA gel formulations to generate soft and stiff gels [60].

#### Collagen functionalization of PA gels

Coverslips were gently removed from the gel surface, and the gels were functionalized with bi-functional crosslinker sulfosuccinimidyl 6-(40-azido-20-nitro- phenylamino) hexanoate (Sulfo-SANPAH; 0.5 mg/mL; ThermoFisher Scientific, 22589) diluted in HEPES buffer (pH 8.5). Gels were then exposed to UV light for 17 min to activate crosslinking and sterilize the gels, followed by three washes with HEPES buffer (pH 8.5). Afterward, gels were treated with collagen I solution (0.2 mg/mL in PBS, pH 7.5) and incubated overnight at 37 °C. The following day, gels were washed 3 times with PBS (pH 7.5) and incubated in INC medium for at least 30 min to equilibrate before seeding HMFs in their relevant medium.

### Human Mammary Fibroblast (HMF) Conditioning with MDA-MB-231 Tumor-Conditioned Medium (mTCM) and Matrix Assembly

To model a cancer-associated fibroblast (CAF)-like phenotype, HMFs were conditioned in mTCM for 3 days, while HMFs conditioned in INC for 3 days were used as the negative control [15, 17]. INC-conditioned HMFs were seeded on soft collagen-functionalized gels (∼2 kPa), whereas mTCM-conditioned HMFs were seeded on stiff collagen-functionalized gels (∼20 kPa) to model the combined biochemical signaling and mechanical cues associated with normal and metastatic breast tissue, respectively. Hereafter, INC-conditioned HMFs cultured on soft collagen-functionalized gels are referred to as the physiological condition, while mTCM-conditioned HMFs cultured on stiff collagen-functionalized gels are referred to as the pathological condition (tumor-associated). HMFs were seeded at a density of 3×10^5^ per 20-mm glass inset. Cells outside of the centrally positioned collagen-functionalized PA gels were not included for downstream imaging and analysis. Physiological and pathological HMFs were allowed to deposit and assemble matrices for 5 days, with the corresponding medium changed on days 1 and 3 after seeding and supplemented with 75 µg/mL of freshly made ascorbic acid to enhance matrix maturation [69–71]. A subset of samples was fixed and immunostained prior to decellularization to assess the combined effect of paracrine signaling and mechanical stiffness of the tumor microenvironment on HMF activation, whereas the remaining samples were decellularized for subsequent macrophage reseeding experiments.

### Decellularization of Conditioned HMF-Derived Matrices

After 5 days of culture, physiological and pathological HMF-derived matrices were decellularized following the Cukierman protocol [63–65]. Briefly, cells were removed using an alkaline detergent (0.5% Triton X-100 and 20 mM NH_4_OH in dPBS), followed by a 5-minute incubation at 37 °C to complete cell lysis. The decellularized matrices were then incubated overnight at 4 °C. The next day, the matrices were washed twice with dPBS and treated with DNase I (10 U/mL; ThermoFisher Scientific, 18047019) in 1X DNase reaction buffer (100 mM Tris–HCl, pH 7.5; 25 mM MgCl□; 1 mM CaCl□) for 30 minutes at 37 °C to eliminate residual DNA debris. Following this, matrices were incubated with 1X DNase inactivation buffer (100 mM Tris–HCl, pH 7.5; free of MgCl□ and CaCl□) for 15 minutes at room temperature, then washed twice more with inactivation buffer. The decellularized matrices were either fixed and immunostained for EDA-FN microarchitectural analysis or stored in dPBS with 1% penicillin–streptomycin at 4 °C for subsequent macrophage reseeding experiments.

### Monocyte-to-Macrophage Differentiation and Reseeding on Decellularized HMF-Derived Matrices

Human peripheral blood monocytes were differentiated into macrophages according to the Spiller protocol [50, 72, 73]. Briefly, human monocytes (1 ×10^7^ per flask) were cultured in Nunc non-treated flasks (ThermoFisher Scientific, 156800) in RPMI medium supplemented with macrophage colony-stimulating factor (M-CSF; 20 ng/mL; PeproTech) for 7 days at 37 °C, with medium changes every 2-3 days. Macrophages were then incubated with TrypLE Select enzyme (ThermoFisher Scientific, 12563011) for 10 min at 37 °C and gently detached using a cell scraper. Before reseeding macrophages, decellularized matrices were equilibrated in RPMI medium for 30 min at 37 °C. Macrophages were then seeded onto physiological and pathological decellularized HMF-derived matrices at a density of 2 × 10^5^ per 20-mm glass inset, cultured for 48 h, followed by fixing and immunostaining.

### Immunofluorescence Staining

#### Immunofluorescence staining of physiological and pathological HMF cultures

Physiological and pathological HMFs were fixed on day 5 using 4% paraformaldehyde (PFA; ThermoFisher Scientific, AAJ61899AK) for 1 h. Samples were then immunostained for DAPI (1:5000; ThermoFisher Scientific, D1306), rabbit polyclonal anti-fibronectin antibody (anti-FN; Sigma-Aldrich, F3648), mouse monoclonal anti-α-smooth muscle actin antibody (α-SMA; 1 µg/mL; ThermoFisher Scientific, 14-9760-82), and Alexa Fluor 568 Phalloidin (anti-F-actin; 1:250; ThermoFisher Scientific, A12380). Corresponding secondary antibodies included goat anti-rabbit Alexa Fluor 488 (1:100; ThermoFisher Scientific, A-11008) and goat anti-mouse Alexa Fluor 647 (1:100; ThermoFisher Scientific, A32728).

#### Immunofluorescence staining of physiological and pathological HMF-derived decellularized matrices

Physiological and pathological HMF-derived matrices were fixed on day 5 and immunostained for DAPI (1:5000; ThermoFisher Scientific, D1306), rabbit polyclonal anti-FN antibody (Sigma-Aldrich, F3648), and mouse monoclonal anti-cellular fibronectin specific for EDA-FN (clone FN-3E2; Sigma-Aldrich, F6140). Appropriate secondary antibodies were used, including goat anti-rabbit Alexa Fluor 488 (1:100; ThermoFisher Scientific, A-11008) and goat anti-mouse Alexa Fluor 647 (1:100; ThermoFisher Scientific, A32728).

#### Immunofluorescence staining of macrophages reseeded on physiological and pathological HMF-derived decellularized matrices

Macrophages reseeded on physiological and pathological HMF-derived matrices were fixed after 48 h. A subset of samples was immunostained for DAPI (1:5000; ThermoFisher Scientific, D1306), mouse monoclonal anti-CD86 antibody (5-30 µg/mL; ThermoFisher Scientific, MA566007), rabbit monoclonal anti-CD206 antibody (1:100; ThermoFisher Scientific, MA5-32498), and Alexa Fluor 568 Phalloidin (anti-F-actin; 1:250; ThermoFisher Scientific, A12380). Corresponding secondary antibodies were used, including goat anti-rabbit Alexa Fluor 488 (1:100; ThermoFisher Scientific, A-11008) and goat anti-mouse Alexa Fluor 647 (1:100; ThermoFisher Scientific, A32728). The second subset of samples was immunostained for DAPI (1:5000; ThermoFisher Scientific, D1306), Alexa Fluor 568 Phalloidin (anti-F-actin; 1:250; ThermoFisher Scientific, A12380), and mouse monoclonal anti-cellular fibronectin specific for EDA-FN (clone FN-3E2; Sigma-Aldrich, F6140) with secondary antibody of goat anti-mouse Alexa Fluor 647 (1:100; ThermoFisher Scientific, A32728).

### Confocal Imaging and Analysis

#### ECM organization and fibroblast cytoskeletal analysis

Confocal images were acquired using an Olympus Fluoview FV1200 confocal microscope with a 30X oil immersion objective, a z-step size of 1 µm, and a pinhole size set to 1 Airy unit. Confocal images in OIF format were imported into ImageJ/Fiji (NIH) for further image processing and analysis. Each fluorescence channel was split, and maximum-intensity projections of z-stacks were generated.

Angle distribution and fiber alignment for FN matrices and cytoskeleton were analyzed using the ImageJ OrientationJ plug-in, as previously described by Franco-Barraza et al. [71]. Briefly, maximum-intensity projection images of fibers were processed using the OrientationJ-Distribution tool. The obtained raw data, including a table of orientation angles and corresponding occurrence values for each detected fiber in the image, were imported into a custom spreadsheet tool, adapted from a macro file previously developed [71]. The mode orientation angle was defined as the angle with the highest occurrence value and was normalized to zero, and all other angles were calculated relative to the mode angle. The angle distribution and the percentage of fibers oriented within + 15° of the mode angle were calculated. For visualization of fiber angle distributions, images were processed using the OrientationJ_Analysis tool. The resulting images were then imported into Adobe Photoshop, and hue values were adjusted to display the mode angle in cyan color. The thickness of physiological and pathological HMF-derived matrices was measured by quantifying the number of total FN-positive z-planes, with the z-step set to 1 µm on the confocal microscope.

The percentage of α-SMA incorporated into F-actin was quantified using an area-based colocalization approach in ImageJ/Fiji. Maximum-intensity projection images of α-SMA and F-actin channels were converted to 8-bit grayscale, and binary masks were generated using the Auto Local Threshold (Phansalkar) method, applied consistently across physiological and pathological conditions. The binary masks were combined using the Image Calculator (AND function), and the area of the resulting overlap mask was measured using the Analyze Particles function and normalized to the α-SMA-positive area. To generate representative images of α-SMA incorporation into F-actin, maximum intensity projection images of α-SMA were converted to 8-bit grayscale. The α-SMA/F-actin overlap mask, generated using the Image Calculator (AND function), was overlaid onto the corresponding α-SMA images using the ImageJ overlay function with fixed transparency.

The JACoP plugin in ImageJ was used for quantifying EDA-FN incorporation into total-FN. Paired maximum intensity projections of EDA-FN and total FN were imported and automatically thresholded by the plugin, and Manders’ colocalization coefficient M2, representing the fraction of EDA-FN overlapping with total FN, was quantified. Higher M2 values indicate greater incorporation of EDA-FN into the total FN network. To quantify EDA-FN and total FN fiber width, CT-FIRE was used, which applies a curvelet-transform (CT) based denoising filter, followed by an automated fiber extraction using the tracking algorithm called fiber extraction (FIRE) [74, 75]. The intrafibrillar space in EDA-FN and total-FN matrices was quantified using a custom-written ImageJ macro. Maximum-intensity projection images of EDA-FN and total FN channels were converted to 8-bit grayscale, and binary masks were generated using the Auto Local Threshold (mean) method, applied consistently across physiological and pathological conditions. The binary images were then inverted to represent intrafibrillar spaces as foreground. Intrafibrillar space areas were then quantified using the Analyze Particle function.

#### Macrophage morphology and phenotypic analysis on EDA-FN matrices

EDA-FN matrix coverage following macrophage reseeding was quantified by background subtraction and application of a uniform thresholding method (Default Threshold) across both physiological and pathological conditions. Binary masks were generated, and the percentage area was quantified using the Color Pixel Counter plugin in ImageJ/Fiji. Fluorescence images of macrophages reseeded on the EDA-FN matrices were acquired using the same confocal microscope mentioned above (30X objective). Individual macrophages were detected using the phalloidin channel and segmented using CellPose, a deep learning-based segmentation algorithm [76]. The resulting masks were then imported into ImageJ/Fiji for morphological analysis, including cell spreading area, circularity, and aspect ratio (Online Resource 1). Super-resolution images of macrophage protrusions interacting with EDA-FN matrices were acquired using a Nikon AXR resonant-scanning confocal with NSPARC spatial array detector on an Eclipse Ti2E inverted microscope with a 60X objective.

Phalloidin-based segmentation masks generated with CellPose were used to quantify the mean fluorescence intensity (MFI) of CD86 and CD206 markers for each cell. Background-subtracted MFI values were calculated for each cell per channel and subtracted from single-cell MFI values. The CD86-to-CD206 corrected MFI ratio was then determined for each macrophage. The average MFI values of each marker were quantified per ROI across at least three independent replicates. To generate a scatter plot, single-cell corrected MFI values for CD86 and CD206 markers were plotted for macrophages seeded on physiological and pathological EDA-FN matrices. The median MFI values of each marker were used to determine a quadrant threshold, classifying macrophages into CD86-dominant, CD206-dominant, and hybrid populations. To quantify the percentage population of macrophages, the number of macrophages classified as CD86-dominant, CD206-dominant, or hybrid in each physiological and pathological condition was normalized to the total number of macrophages per condition.

### Statistical Analysis

All experiments used a minimum of at least three samples, with analysis of at least four regions of interest per sample. GraphPad Prism was used for all statistical analyses. Comparison between two groups was performed using the Mann-Whitney U test for non-normally distributed data and the Welch’s t-test for normally distributed data. Comparison between two independent variables was performed using two-way ANOVA, followed by post-hoc Tukey’s multiple comparisons test.

## Results

### HMF-derived Total FN Matrix Assembly and Cytoskeletal Organization Prior to Decellularization

To assess the combined impact of tumor-conditioned factors, representing paracrine signaling from an aggressive breast cancer, and substrate stiffness on HMF phenotype, HMFs were cultured under physiological (INC medium + soft PA gel) or pathological (mTCM medium + stiff PA gel) conditions and fixed. Pathological HMFs assembled highly aligned FN matrices compared to physiological HMFs (Fig. 2A). Pathological FN matrices had a narrower averaged angle distribution normalized to the mode angle (Fig. 2B), accompanied by a significantly higher percentage of FN fibers oriented within +15° of the mode angle (Fig. 2C). Also, hue-normalized orientation map images revealed that, under pathological conditions, FN matrices were predominantly oriented toward the mode angle (shown in cyan), whereas physiological FN matrices displayed disorganized orientation angles (Fig. 2D).

**Fig. 2.**
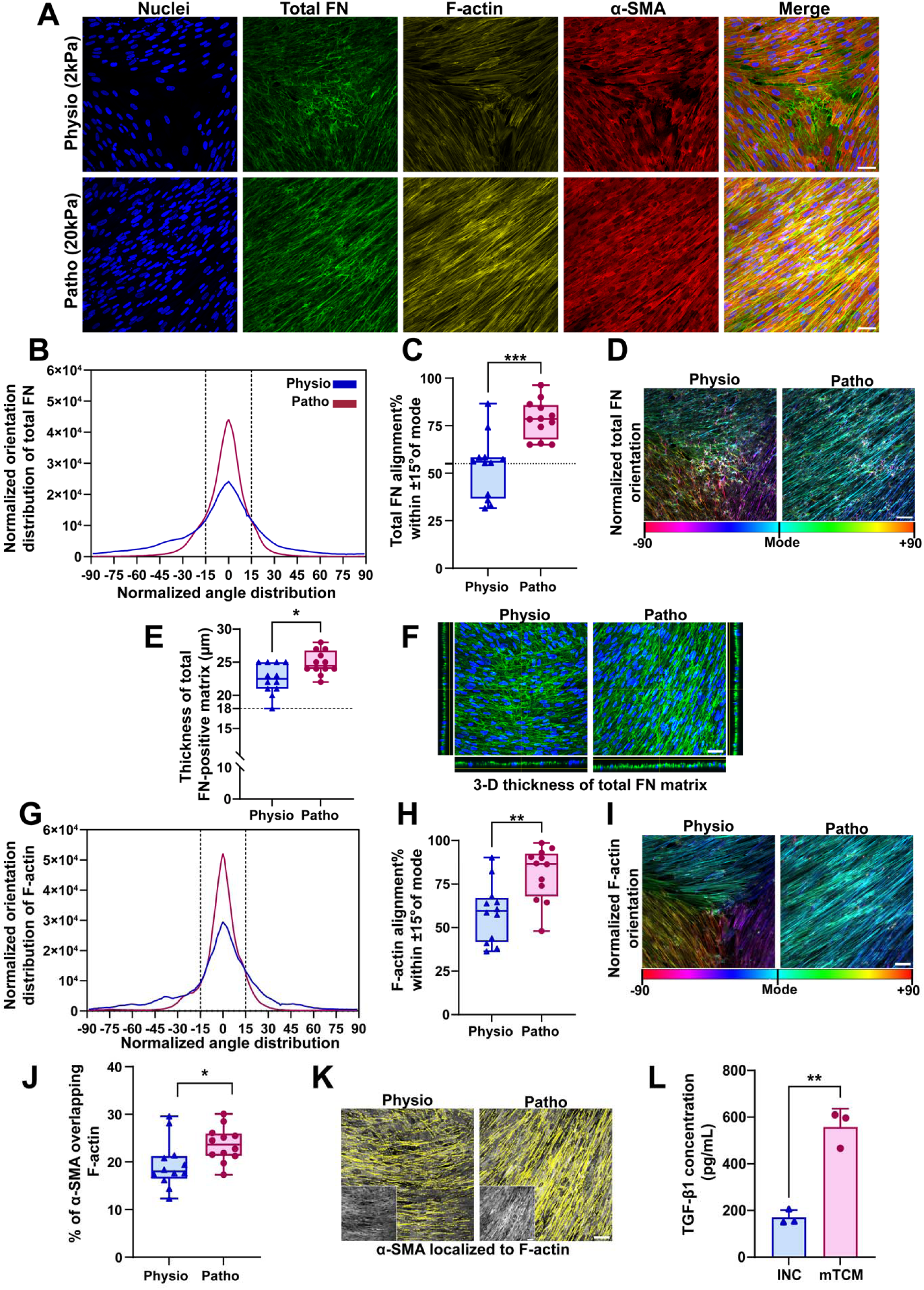
Pathological HMFs exhibit altered FN matrix assembly and cytoskeletal organization. **A** Representative confocal microscopy images of physiological and pathological HMFs and associated fibronectin matrices. Nuclei are shown in blue, total FN in green, F-actin in yellow, and α-SMA in red, and the merged image of all four channels. Scale bar, 50 µm. **B** Averaged variations of angle distributions of total FN matrices normalized to 0° mode. **C** Percentage of total FN matrices within 15° of the mode angle. The dotted line denotes 55% alignment. Each data point represents the mean value per region of interest (ROI) from at least 3 independent replicates. **D** The corresponding total FN matrix angle distributions, determined using ImageJ’s *Orientation J* plugin, were normalized using hue values in Photoshop to visualize the mode angle as cyan. Scale bar, 50 µm. **E** Semi-quantitative measurement of total FN matrix thickness determined by total FN-positive z-planes, with the z-step size of 1 μm during confocal imaging. **F** Representative merged confocal microscopy images of total FN-positive matrices and nuclei. XZ and YZ orthogonal views are shown. Scale bar, 50 µm. **G** Averaged variations of angle distributions of F-actin fibers normalized to 0° mode. **H** Percentage of F-actin within 15° of the mode angle. **I** The corresponding F-actin angle distributions, determined using ImageJ’s *Orientation J* plugin, were normalized using hue values in Photoshop to visualize the mode angle as cyan. Scale bar, 50 µm. **J** Percentage of α-SMA-positive area incorporated into F-actin. Each data point represents the mean value per ROI from at least 3 independent replicates. **K** Representative images of a mask of α-SMA-positive area incorporated into the F-actin, shown in yellow, overlaid on raw α-SMA images, shown in grey. Scale bar, 50 µm. **L** TGF-β1 concentration (pg/mL) in mTCM-and INC-conditioned media measured by ELISA. Each data point represents the mean of an independent experiment (n = 3), with three technical replicates averaged. Data is shown as median ± interquartile range. Asterisks indicate * *p* < 0.05, ** *p* < 0.01, *** *p* < 0.001, **** *p* <0.0001. Physio refers to INC-conditioned HMFs cultured on soft collagen-functionalized gels, and patho refers to mTCM-conditioned HMFs cultured on stiff collagen-functionalized gels

Both physiological and pathological HMF assembled total FN matrices of > 18 µm. Under pathological conditions, total FN matrices were thicker compared to physiological controls (Fig. 2E). Increased total FN deposition under pathological conditions is further shown by orthogonal XZ and YZ views in Fig. 2F.

Under pathological conditions, HMFs exhibited altered cytoskeletal organization (Fig. 2A). Pathological HMFs assembled highly aligned F-actin fibers compared to physiological controls, as displayed by a tighter average angle distribution (Fig. 2G) and a significant increase in the percentage of F-actin fibers aligned within + 15° of the mode angle (Fig. 2H). The hue-normalized orientation map images of F-actin fibers suggest that F-actin fibers were organized co-aligned with FN matrices under both physiological and pathological conditions (Fig. 2I, D).

To investigate HMF activation, we quantified the percentage of α-SMA-positive area incorporated into F-actin stress fibers. Pathological HMFs exhibited higher incorporation of α-SMA into F-actin stress fibers compared to physiological controls, suggesting increased HMF activation under pathological conditions (Fig. 2J). This was further displayed by overlaying a yellow mask representing α-SMA-F-actin overlap onto representative grayscale α-SMA images, which qualitatively indicates the extent of α-SMA incorporation into F-actin stress fibers under physiological and pathological conditions (Fig. 2K). To evaluate what might be contributing to these pathological changes in cytoskeletal and matrix organization, looked at TGF-β1. Measurement of TGF-β1 concentration in mTCM and INC media revealed an approximate threefold increase in mTCM compared to INC (Fig. 2L)

### EDA-FN and Total FN Matrix Organization After Decellularization

To develop a cell-free ECM platform for studying EDA-FN-macrophage interactions, HMF-derived matrices were decellularized. The organization of EDA-FN and total FN was analyzed under both physiological and pathological conditions. The absence of DAPI staining confirms successful decellularization while maintaining matrix organization, allowing direct analysis of total FN and EDA-FN organization independent of cellular contributions (Fig. 3A). Quantitative colocalization analysis showed a significantly higher Mander’s overlap coefficient (M_2_) between EDA-FN and total FN under pathological conditions compared to physiological controls, indicating increased integration of EDA-FN into total FN fibers (Fig. 3B). OrientationJ analysis revealed notable differences in the matrix alignment of total FN and EDA-FN between physiological and pathological conditions (Fig. 3C-G). Pathological total FN matrices displayed a narrower average angle distribution relative to physiological controls, suggesting enhanced fiber alignment (Fig. 3C). Additionally, hue-normalized orientation map images demonstrated that pathological total FN matrices were mainly aligned toward the mode angle (shown in cyan), whereas physiological control matrices were oriented toward multiple angles in a disorganized manner (Fig. 3D).

**Fig. 3.**
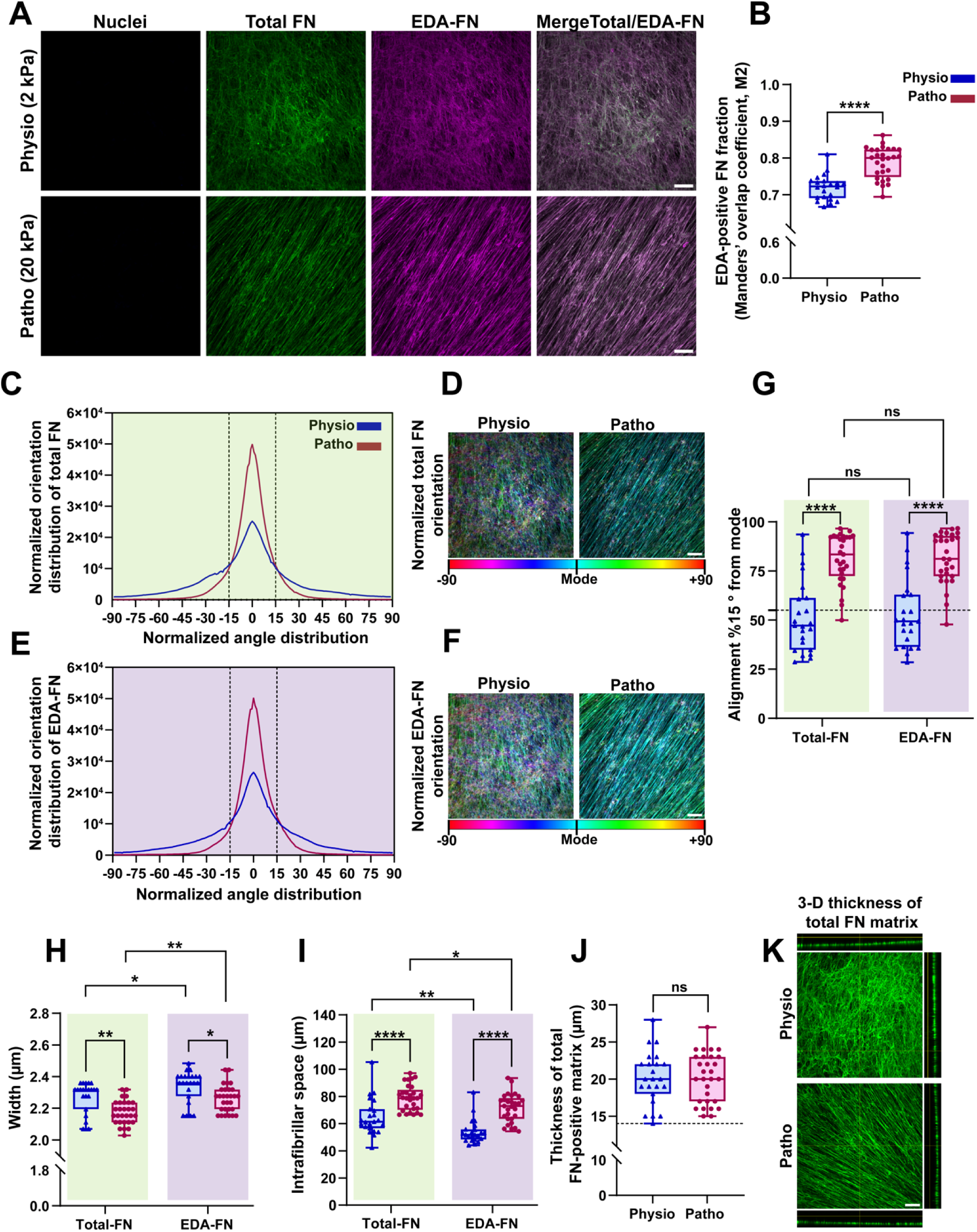
Decellularized pathological EDA-FN matrices have aberrant organization. **A** Representative confocal microscopy images of physiological and pathological decellularized total FN and EDA-FN matrices. Nuclei are shown in blue, total FN in green, EDA-FN in magenta, and the merged image of total FN and EDA-FN. Scale bar, 50 µm. **B** Mander’s colocalization coefficient (M2), quantifying the fraction of EDA-FN overlapping with total-FN, was quantified using the JACoP plugin in ImageJ. Each data point represents the mean value per ROI from at least 3 independent replicates. **C** Averaged variations of angle distributions of total FN matrices normalized to 0° mode. **D** The corresponding total FN matrix angle distributions, determined using ImageJ’s *Orientation J* plugin, were normalized using hue values in Photoshop to visualize the mode angle as cyan. Scale bar, 50 µm. **E** Averaged variations of angle distributions of EDA-FN matrices normalized to 0° mode. **F** The corresponding EDA-FN matrices’ angle distributions, determined using ImageJ’s *Orientation J* plugin, were normalized using hue values in Photoshop to visualize the mode angle as cyan. Scale bar, 50 µm. **G** Percentage of total FN and EDA-FN matrices within 15° of the mode angle. The dotted line denotes 55% alignment. **H** Fiber width of decellularized total FN and EDA-FN, quantified by CT-FIRE. **I** Intrafibrillar space of decellularized total FN and EDA-FN matrices, calculated using ImageJ. **J** Semi-quantitative measurement of decellularized total FN matrix thickness determined by total FN-positive z-planes, with the z-step size of 1 μm during confocal imaging. **K** Representative confocal microscopy images of decellularized total FN-positive matrices. XZ and YZ orthogonal views are shown. Scale bar, 50 µm. Data is shown as median ± interquartile range. Asterisks indicate * *p* < 0.05, ** *p* < 0.01, *** *p* < 0.001, **** *p* < 0.000

Similarly, pathological EDA-FN matrices exhibited a narrower averaged angle distribution compared to physiological controls (Fig. 3E). This was further supported by hue-normalized orientation map images of pathological EDA-FN matrices, in which fibers were predominantly aligned toward the mode angle (Fig. 3F). Visual observation of hue-normalized orientation maps of total FN and EDA-FN suggests that fibers from both matrices were co-aligned toward similar directions under both physiological and pathological conditions. Consistent with the orientation distribution analysis, pathological total FN and EDA-FN matrices both exhibited a significantly higher percentage alignment compared to physiological controls. However, no statistically significant difference was detected between the alignment of total FN and EDA-FN within either condition, suggesting that the observed differences in alignment were driven by whether matrices were developed under physiological or pathological conditions rather than FN isoform (Fig. 3G).

Additional quantitative characterization of total FN and EDA-FN matrices revealed both condition-dependent and isoform-dependent microarchitectural alterations in matrices. Under pathological conditions, both total FN and EDA-FN matrices displayed a significant decrease in fiber width compared to their physiological controls (Fig. 3H). Notably, under both physiological and pathological conditions, EDA-FN matrices exhibited a thicker fiber width than that of total FN matrices (Fig. 3H).

In contrast to fiber width, intrafibrillar space was significantly increased in pathological matrices for both total FN and EDA-FN matrices respective to physiological controls (Fig. 3I). Also, under both conditions, EDA-FN matrices exhibited reduced intrafibrillar space than total FN matrices (Fig. 3I). Collectively, these findings show a negative correlation between fiber width and intrafibrillar space for both total FN and EDA-FN matrices, with pathological matrices displaying thinner fibers and enlarged intrafibrillar spaces (Fig. 3H, I).

Semi-quantitative thickness analysis revealed no statistically significant difference in decellularized total FN matrix thickness between physiological and pathological conditions (Fig. 3J). However, the decellularization process resulted in a relative reduction in total FN matrix thickness, decreasing from > 18 µm prior to decellularization to > 14 µm (Fig. 2E, 3J).

### Macrophage Morphology and Remodeling of HMF-Derived EDA-FN Matrices

To investigate the impact of pathological HMF-derived EDA-FN-rich matrices on macrophage morphology and their interaction with said matrix, macrophages were seeded onto physiological and pathological matrices. As indicated by yellow arrows (Fig. 4A), pathological HMF-derived matrices exhibit localized loss of EDA-FN signal beneath the macrophage cell body, forming perforations, along with fiber bundling around the cell body, whereas physiological matrices show a more diffuse loss of signal throughout. These patterns suggest active matrix remodeling by macrophages. Consistent with this, quantification of EDA-FN matrix coverage following macrophage reseeding revealed higher pathological matrix coverage area compared to that of physiological matrices (Fig. 4B). Macrophage morphology was analyzed across conditions. There were no statistically significant differences in cell spreading area between physiological and pathological matrices (Fig. 4C). Although macrophages displayed heterogeneous morphologies, including rounded, star-shaped, and elongated forms on both matrices, macrophages attached on pathological HMF-derived EDA-FN matrices were associated with higher circularity and lower aspect ratio compared to macrophages attached onto physiological HMF-derived EDA-FN matrices (Fig. 4D, E).

**Fig. 4.**
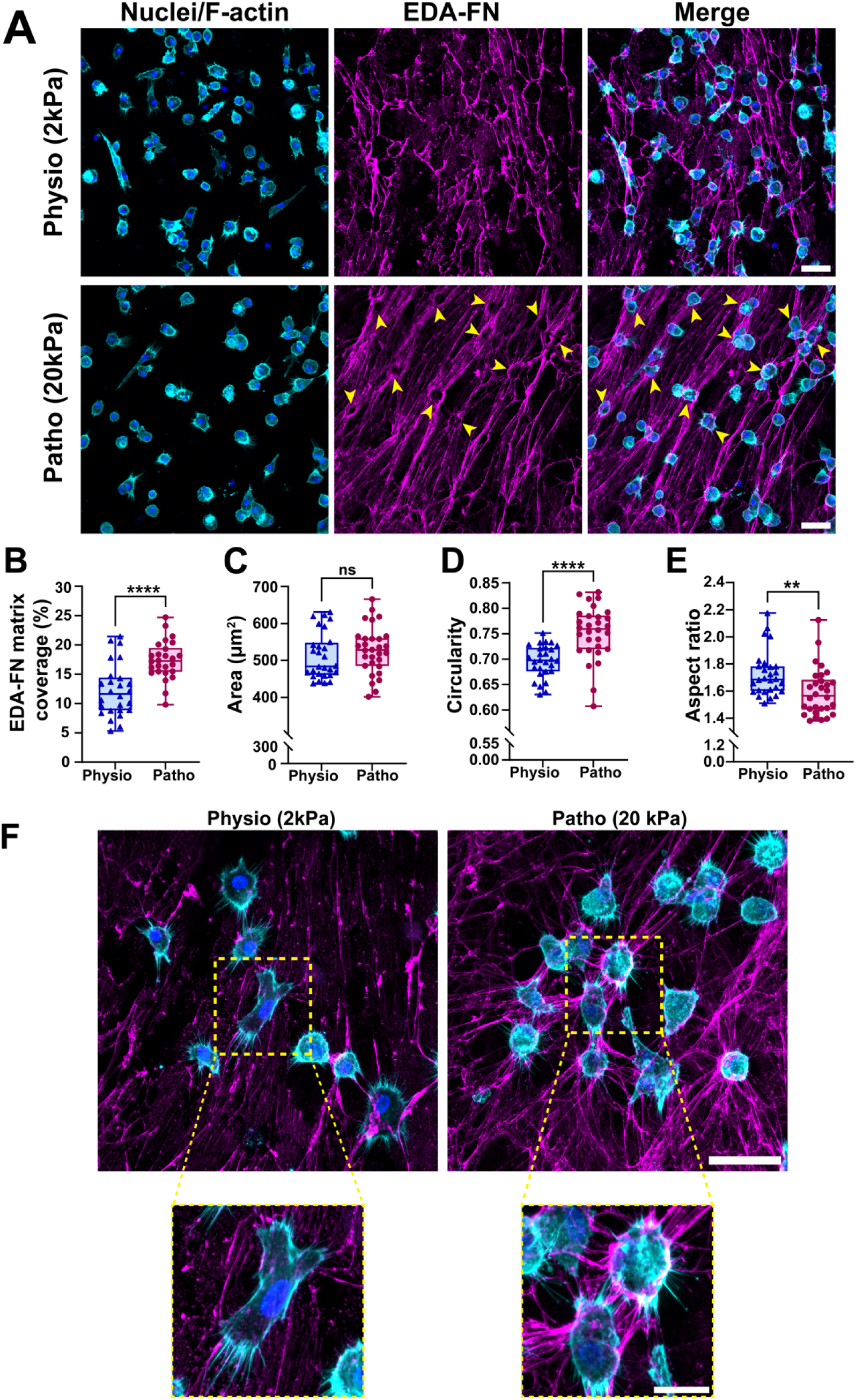
Macrophages display heterogeneous morphologies on HMF-derived EDA-FN-rich matrices and are associated with EDA-FN matrix reorganization. **A** Representative confocal microscopy images of macrophages reseeded on physiological and pathological HMF-derived EDA-FN matrices. Nuclei are shown in blue, F-actin in cyan, EDA-FN in magenta, along with a merged image of all channels. Yellow arrows indicate localized regions of reduced EDA-FN signal beneath macrophages on pathological HMF-derived EDA-FN matrices. Scale bar, 50 µm **B** Quantification of EDA-FN matrix area coverage following macrophage reseeding. Quantification of macrophage morphology, including **C** spreading area, **D** circularity, **E** and aspect ratio. Each data point represents the mean value per ROI from at least 3 independent replicates. **F** Super-resolution microscopy images display macrophage protrusions interacting with EDA-FN fibers. Scale bar, 50 µm; zoomed images, 10 µm. Data is shown as median ± interquartile range. Asterisks indicate * *p* < 0.05, ** *p* < 0.01, *** *p* < 0.001, **** *p* < 0.0001

Super-resolution microscopy images revealed macrophage protrusions close to EDA-FN fibers, suggesting cell-matrix interactions (Fig. 4F). Zoomed-in images showed the cell-matrix interface, where macrophage protrusions were extended into the surrounding EDA-FN matrix. In pathological HMF-derived EDA-FN matrices, fibers were more locally clustered and reorganized around macrophages, consistent with localized EDA-FN signal loss and higher matrix coverage in pathological matrices following macrophage reseeding (Fig. 4A, B, F).

### Macrophage Phenotype on HMF-Derived EDA-FN Matrices

Immunostaining for CD86 and CD206, pro-inflammatory and anti-inflammatory markers, respectively, in macrophages reseeded on physiological and pathological HMF-derived EDA-FN matrices revealed that cells were predominantly double-positive for both markers (Fig. 5A). However, macrophages attached on pathological HMF-derived EDA-FN matrices showed slightly higher CD206 intensity, whereas physiological matrices displayed a higher CD86 intensity. Consistent with this, the CD86-to-CD206 MFI ratio was decreased in macrophages on pathological matrices compared to macrophages on physiological matrices, suggesting a shift toward CD206 on pathological matrices (Fig. 5B). Single-cell analysis of CD86 and CD206 MFI indicated that macrophages were predominantly distributed diagonally, suggesting co-expression of both markers (hybrid population) with smaller subsets categorized as CD86-dominant (CD86^+^) or CD206-dominant (CD206^+^) (Fig. 5C). Using median-based threshold quadrants, pathological matrices were associated with a slight shift of macrophages toward a CD206-dominant population, with a corresponding reduction in CD86-dominant cells. This is then visualized as a population percentage, where macrophages were primarily categorized into the hybrid population, with a shift toward CD206-dominant cells on pathological matrices (Fig. 5D).

**Fig. 5.**
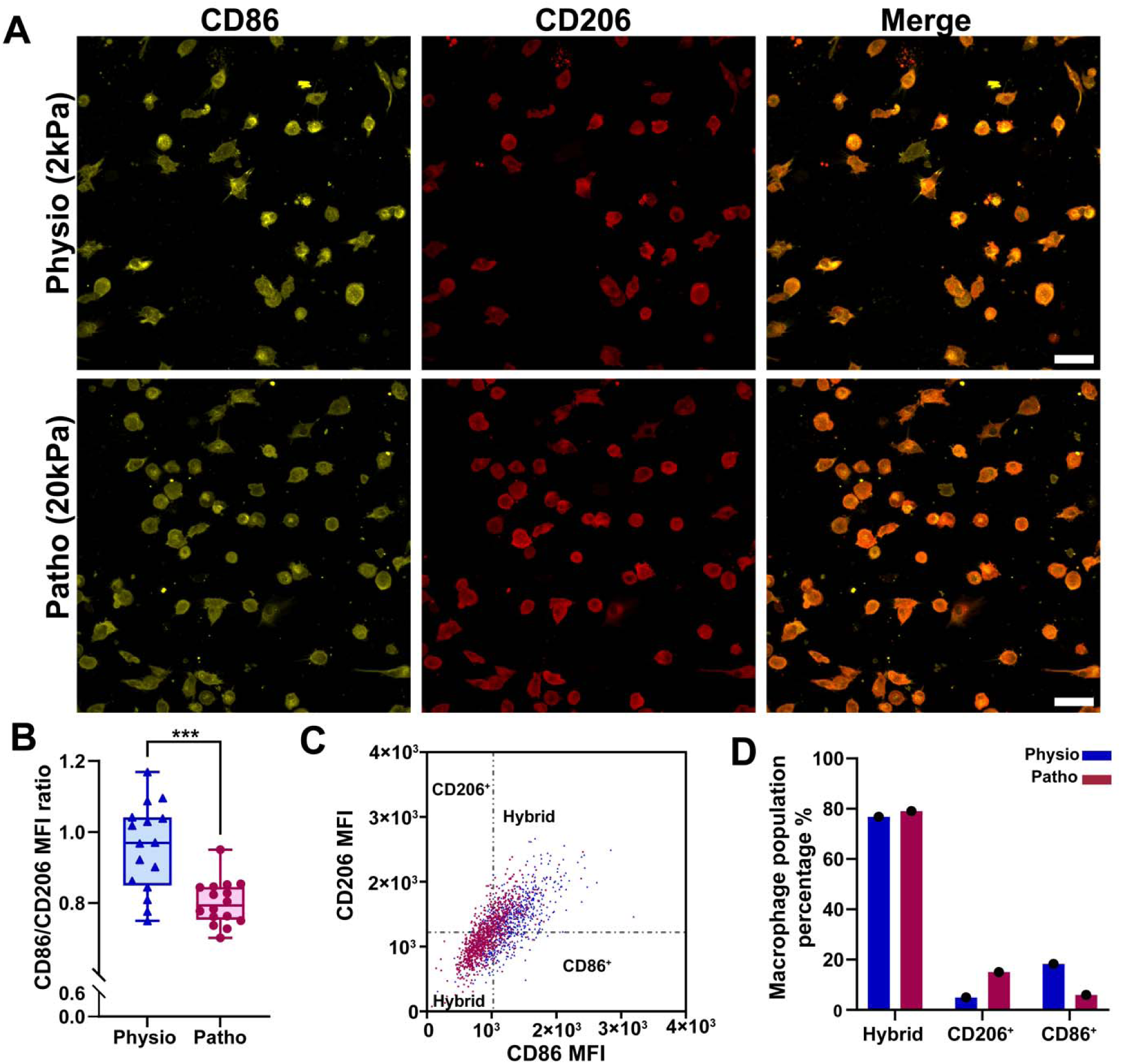
Macrophages adopt a predominantly hybrid phenotype on HMF-derived EDA-FN matrices, with macrophages on pathological HMF-derived EDA-FN matrices associated with increased CD206 signal intensity. **(A)** Representative confocal microscopy images of macrophages cultured on physiological and pathological EDA-FN matrices. Nuclei are shown in blue, CD86 in yellow, CD206 in red, and a merged image of all channels. **(B)** Ratio of CD86 to CD206 mean fluorescence intensity (MFI). Each data point represents the mean value per ROI from independent replicates. **(C)** Single-cell scatter plot of CD86 and CD206 MFI in macrophages. Each data point represents one cell. Dashed lines show threshold-based quadrant separation based on the median MFI of each marker **(D).** Quantification of macrophage population categorized as hybrid, CD206-dominant (CD206^+^), or CD86-dominant (CD86^+^) based on MFI thresholds. Asterisks indicate * *p* < 0.05, ** *p* < 0.01, *** *p* < 0.001, **** *p* < 0.0001

## Discussion

In this study, we developed a human mammary fibroblast-derived matrix model that mimicked tumor-associated biochemical and mechanical cues within the tumor stroma microenvironment to study matrix-specific effects on macrophage behavior. Using this platform, we found that pathological biochemical and mechanical cues induced HMF activation to a CAF-like phenotype, which was accompanied by cytoskeletal reorganization and significant microarchitectural alterations in total FN matrices. Following decellularization, the matrices preserved their structural organization, with both pathological total FN and EDA-FN displaying aberrant microarchitectural characteristics. Collectively, this platform establishes a robust ECM network model, allowing the investigation of macrophage interactions with tumor-associated EDA-FN-rich matrices.

The dynamic interplay between aberrant biochemical signaling and mechanical cues distinguishes the tumor microenvironment from normal tissue, promoting fibroblast activation and subsequent pathological ECM remodeling [36, 77–79]. Tumor-derived soluble factors, particularly TGF-ß1 signaling, are well-established drivers of fibroblast activation and differentiation into CAFs within the tumor microenvironment [80–82]. α-SMA, a known activated fibroblast marker, is incorporated into actin stress fibers under increased tension and contributes to fibroblast contractility [83, 84]. Consistent with this, the mTCM in our system contained elevated levels of TGF-ß1 compared to INC. Additionally, we observed enhanced incorporation of α-SMA into F-actin stress fibers in fibroblasts under pathological conditions (mTCM + stiff substrate). It is important to note that the presence of α-SMA in fibroblasts under physiological conditions could be explained by the fact that, before re-seeding onto soft PA gels and switching to low-serum media, the cells were cultured on plastic tissue culture flasks and fed high-serum media. Such exposure to the combination of stiff plastic tissue culture and latent TGF-ß present in serum may trigger partial activation of fibroblasts [27, 85, 86]. This is consistent with previous studies showing fibroblasts can retain memory of previous exposure to biochemical and mechanical cues [87]. Together, our findings suggest that the combined biochemical and mechanical cues in our model can drive fibroblast activation into a CAF-like phenotype [88, 89].

Activated fibroblasts deposit and assemble abnormally organized ECM proteins such as FN and collagen, resulting in a stiffer tumor tissue [12, 28, 79]. In turn, enhanced tissue stiffness further activates fibroblasts, creating a continuous feedforward loop maintaining CAF phenotype [32, 80]. Activated fibroblasts reorganize FN into highly aligned, parallel fibers through actomyosin-mediated contractility and traction force generation [81, 82]. In line with this, we observed that under pathological conditions, HMFs assembled highly organized, aligned FN fibers, accompanied by increased F-actin stress fiber alignment compared to physiological conditions. Notably, FN fibers and F-actin stress fibers were co-aligned. This suggests coordinated reorganization of the ECM and the intracellular cytoskeleton. Previous studies have shown that FN fiber orientation is determined by the orientation of matrix assembly sites, which are linked to focal adhesions and actin stress fibers [83]. And aligned ECM has been shown to promote the formation of highly aligned actin stress fibers in fibroblasts, which govern directed cell migration [84]. Together, these findings suggest bidirectional reorganization between ECM and cytoskeleton architecture. Our findings further demonstrate that pathological conditions shift this coordinated extracellular and intracellular organization toward a highly aligned architecture, a feature characteristic of CAF-like phenotype.

The ECM is a complex microenvironment composed of fibrillar proteins and soluble factors that not only provide cells with structural support but also regulate their functional behavior. Conventional 2D and simplified 3D systems, such as glass, plastic, pure collagen or FN hydrogels, do not fully mimic the physiological microenvironment [85, 86]. However, cell-derived matrices provide a more in vivo-like microenvironment and allow us to study cell-ECM interactions and underlying signaling mechanisms [87, 88]. The decellularization process enables the removal of cellular components while preserving matrix composition, fibrillar architecture, and sequestered growth factors. Accordingly, in our present model, following decellularization, cellular components were successfully removed while preserving overall matrix organization. This pathological cell-derived FN matrix model provides a platform to investigate how tumor-associated FN-rich matrices regulate macrophage behavior by isolating the matrix-derived effects from ongoing soluble factor signaling.

EDA-FN, an alternatively spliced variant of cellular FN, is predominantly deposited in tumor stroma by CAFs and has been linked to fibroblast activation, fibrosis, inflammatory responses, cancer progression, and poor patient prognosis [89, 90]. Furthermore, EDA-FN has been shown to promote FN fibrilogenesis through enhancing FN assembly and stress fiber formation in fibroblasts [91, 92]. Total FN has been shown to colocalize with EDA-FN in head and neck tumor tissues [31]. In accordance with this, we observed an enhanced incorporation of EDA-FN into the fibrillar FN network under pathological conditions, suggesting increased EDA-FN integration in tumor-associated matrices. This elevated incorporation of EDA-FN into the FN network under pathological conditions further supports the tumor-relevant nature of our cell-derived matrix model to study how tumor-associated EDA-FN organization influences surrounding cell behavior.

The ECM is a crucial dynamic component of TME that undergoes substantial remodeling during breast cancer progression [14, 36]. ECM remodeling involves alterations in not only matrix composition but also in microstructural organization, such as alignment and spatial distribution, which in turn modulate cellular functions. ECM alignment is a hallmark of tumor invasion and has been identified as a prognostic marker in breast cancer carcinoma, where aligned fibrillar collagen fibers are associated with poor clinical outcomes [93]. In addition, CAFs have been shown to generate highly aligned FN and collagen matrices that promote directional cancer cell migration [82, 94, 95]. These findings underscore ECM alignment as a key feature of tumor-associated matrices. In our decellularized matrices, pathological conditions were associated with greater alignment of both total FN and EDA-FN compared to physiological conditions. Notably, no difference was observed between total FN and EDA-FN within each condition, suggesting that matrix alignment is primarily governed by pathological condition rather than FN isoform composition. Additionally, the co-alignment of EDA-FN and total FN suggests that EDA-FN fibers are incorporated into the overall FN network along the same orientation axis rather than forming a distinct directional pattern. Altogether, these findings suggest that tumor-associated biochemical and mechanical cues generate a highly organized and aligned FN matrix, in which EDA-FN fibers are co-aligned and integrated within the same fibrillar network. These findings are also consistent with the fibroblast-mediated matrix organization before decellularization, showing that the cell-derived matrix remodeling is maintained following decellularization.

In our platform, pathological conditions were also associated with structural alterations at smaller length scales, where both total FN and EDA-FN exhibited reduced fiber width and enhanced intrafibrillar spacing compared to physiological conditions. In addition to condition-dependent reorganization, isoform-specific differences were observed, with EDA-FN matrices having greater fiber width and lower intrafibrillar spacing compared to total FN under physiological and pathological conditions. This suggests that despite co-alignment, EDA-FN exhibits a structurally distinct organization at smaller length scales within the overall total FN network, highlighting isoform-specific microarchitectural features of the current platform [106].

TAMs are among the most prevalent inflammatory stromal cells within the TME, and their increased infiltration within tumors is often associated with poor clinical outcomes [96, 97]. TAMs play a pivotal role in enhancing tumor invasiveness and metastasis by contributing to ECM remodeling and angiogenesis [40, 98–102]. In addition to soluble tumor-derived factors, mounting evidence suggests that the composition and physical characteristics of the TME can regulate macrophage function, including adhesion, morphology, phenotype, and migration, through cell-matrix interactions and mechanotransduction signaling [37, 46, 48, 103]. Collectively, these findings highlight the significance of ECM organization in modulating macrophage behavior.

Consistent with the increased incorporation of EDA-FN within the FN network observed in our tumor-associated decellularized platform, TAMs have been frequently detected in EDA-FN-rich regions of triple-negative breast tumors [104]. In addition, EDA-FN has been associated with inflammatory responses within the TME, suggesting a spatial and functional association between EDA-FN and TAMs [19, 55, 57, 105]. These findings underscore a prominent role for cell-matrix interactions in regulating TAMs. In this context, our tumor-associated EDA-FN-rich cell-derived matrix platform with distinct organization enables investigation of how EDA-FN organization modulates macrophage function independent of soluble signaling.

Macrophages have been shown to play a prominent role in ECM remodeling within the TME through secretion of matrix metalloproteinases (MMPs), degradation of matrix components, and modulation of ECM organization, thereby supporting tumor invasion [47, 48, 51, 117]. However, these studies have been largely focused on collagen, which further highlights the novelty of our tumor-associated EDA-FN-rich cell-derived matrices. In our decellularized matrices, we observed that, following macrophage reseeding, EDA-FN matrix coverage remained higher under pathological conditions compared to physiological conditions. In line with previous studies showing that macrophage-mediated degradation can occur in localized areas beneath the cell body [118–120], the presence of localized perforations and fiber clustering and bundling around the macrophage cell body in pathological matrices suggests active matrix remodeling, potentially involving both proteolytic activity and mechanical force generation [121]. This highlights the need to further determine the underlying mechanisms by which macrophages degrade and remodel the surrounding matrix. In addition, the presence of finger-like long protrusions extending along EDA-FN fibers at the cell-matrix interface further suggests that macrophages actively interact with and sense the EDA-FN fibrillar network.

Collectively, these findings show that macrophages reorganize EDA-FN matrices and suggest tumor-associated matrix-derived cues can influence how macrophages reorganize the surrounding matrix.

Macrophages sense and respond to mechanical cues from the ECM through integrins, forming focal adhesion-like structures at the cell-matrix interface and connecting the extracellular cues to the intracellular cytoskeleton [45, 46, 122, 123]. This enables bidirectional matrix-cell signaling. Cytoskeletal remodeling of macrophages in response to microarchitectural alterations of the surrounding environment has been shown to regulate cell shape, phenotype, and contribution to fibrotic remodeling [45, 112, 113]. This highlights the significance of matrix-derived cues in macrophage functional behavior. In our platform, macrophages exhibited heterogeneous morphological characteristics across both pathological and physiological ED-FN-rich matrices, with the majority of macrophages displaying a spread, rounded morphology with long protrusions. Notably, pathological matrices were associated with a shift toward lower aspect ratios and higher circularity, despite no statistically significant difference in cell spreading area between conditions. This suggests that matrix-derived cues regulate macrophage morphology independent of cell spreading.

In line with previous studies showing that the mechanical and adhesive properties of macrophage substrate regulate their cytoskeleton-dependent protrusive activity and morphology [125], heterogeneous protrusive patterns were observed in our EDA-FN-rich platforms across conditions. Protrusive rounded cells often exhibited long protrusions distributed symmetrically around the cell body and along surrounding EDA-FN fibers, whereas elongated macrophages were aligned along EDA-FN fibers with protrusions distributed more asymmetrically. These differences in macrophage protrusive patterns suggest that macrophages engage with the matrix in structurally distinct manners, which may reflect underlying cytoskeleton-dependent organization influenced by matrix-derived cues. This is further supported by studies indicating that matrix-derived cues, including organization, composition, and stiffness, regulate macrophage morphological heterogeneity, cytoskeletal organization, phenotype, and migration mode [55, 126]. Collectively, the observed morphological heterogeneity of macrophages, together with the shift toward a more rounded morphology on pathological matrices, suggests that matrix-derived cues play a prominent role in how macrophages interact and structurally adapt to the surrounding matrix via cytoskeleton-dependent alterations in cell morphology.

Macrophages display remarkable phenotypic plasticity within TME and transition between distinct functional states in response to mechanical and biochemical cues received from their surrounding microenvironment [40, 41, 127]. Accordingly, conventional classification of macrophages into classically activated (M1) and alternatively activated (M2) phenotypes is oversimplified and does not fully capture the spectrum of macrophage functional states that exist between these two extremes [43, 128]. The mannose receptor, also known as CD206, is highly expressed on TAMs and is one of the most commonly used markers for macrophages associated with anti-inflammatory, tumor-promoting, and tissue remodeling phenotypes [46, 129, 130]. CD86, however, is a T-cell costimulatory molecule, associated with pro-inflammatory, tumor-suppressing macrophage activity [131–133]. Accordingly, due to the significance of immune evasion and inflammation in cancer progression [40], and the complexity of macrophage phenotypes, in this study, macrophage states are interpreted based on their marker-associated phenotypic profile, where CD206-dominant (CD206^+^) profiles are associated with an anti-inflammatory, pro-tumor-like activity, whereas CD86-dominant (CD86^+^) macrophages are associated with a pro-inflammatory, anti-tumor-like activity.

In our platform, macrophages on EDA-FN- rich matrices co-expressed CD206 and CD86 markers across both conditions, with a large population of macrophages displaying a hybrid phenotype. This is consistent with previous studies showing that tumor-associated cues do not drive macrophages into simplified M1/M2 phenotypes. [109, 134]. Similarly, a recent study challenges the simplified M1/M2 categories, as well as the hypothesis that macrophage phenotypes exist along a spectrum, by showing that within the breast tumor microenvironment, macrophages co-expressed both M1- and M2-associated genes within the same cells, and these genes positively correlated with one another [135].

Macrophages on pathological matrices showed a trend toward a CD206-dominant state, whereas those on physiological matrices showed a trend toward a more CD86-dominant state. This is consistent with previous studies demonstrating that TAMs can co-express markers associated with both anti-inflammatory and pro-inflammatory states, including CD206 and CD86 at the single-cell level [43, 130, 136]. Despite the highly heterogeneous nature of TAMs within tumors, increased prevalence of CD206-positive macrophages has been associated with highly aggressive tumors with higher levels of matrix remodeling, whereas macrophages with pro-inflammatory characteristics were associated with less aggressive tumors [137, 138]. Consistent with this, in our platform, although a large population of macrophages exhibited a hybrid phenotype, macrophages engaging with pathological EDA-FN-rich matrices showed a trend toward a CD206-dominant profile, whereas those on physiological matrices showed a trend toward a more CD86-dominant profile. Collectively, these findings suggest that tumor-associated ED-FN-rich matrices promote macrophage phenotypic heterogeneity. In addition, the observed dysregulated microarchitectural alterations in pathological EDA-FN-rich matrices may contribute to a bias toward the CD206-associated profiles. This suggests that tumor-associated matrix-derived cues can have a significant role in regulating macrophage behavior independent of soluble tumor factors.

In summary, we developed a tumor-associated HMF-derived EDA-FN-rich matrix platform recapitulating dysregulated matrix organization, which has a significant role in breast tumor progression. Following decellularization, this platform eliminated ongoing soluble factor signaling while maintaining distinct organization, enabling us to study how matrix-derived cues shape macrophage behavior. Using this cell-derived platform, we demonstrated that tumor-associated HMF-derived EDA-FN-rich matrices with altered organization can regulate macrophage behavior by promoting cytoskeleton-dependent heterogenenic morphologies and hybrid phenotypic profiles. Given the observed distinct localized loss of EDA-FN signal beneath the macrophage cell body on pathological HMF-derived EDA-FN-rich matrices, and the close interaction between actin-rich protrusions and EDA-FN fibers at the cell-matrix interface, these findings suggest that macrophages are likely to play a key role in remodeling the surrounding EDA-FN matrix under tumor-associated matrix-driven cues. Furthermore, it is important to note that in the present study, we isolated the impact of tumor-associated HMF-derived EDA-FN-rich matrices from tumor-derived soluble factors that macrophages would encounter in vivo. Therefore, future work should incorporate tumor-derived soluble factors into this platform to study how macrophages remodel and degrade the surrounding EDA-FN-rich matrices.

## Conclusion

In this study, we developed a tumor-associated HMF-derived matrix model by exposing human mammary fibroblasts (HMFs) to the combined impact of tumor-derived soluble factors and pathological substrate stiffness to capture the structural imprint of tumor-associated biochemical and mechanical cues within the matrix. These conditions induced a CAF-like phenotype and drove a distinct organization of assembled FN matrices enriched in cellular EDA-FN. Decellularization enabled us to isolate the impact of matrix-specific effects to investigate how tumor-associated HMF-derived EDA-FN-rich matrices regulate macrophage behavior. We showed that there is a bidirectional relationship between macrophages and HMF-derived EDA-FN-rich matrices, where matrix-specific cues regulate macrophage morphology and phenotype. These macrophages actively interacted with the matrix through protrusive cytoskeletal structures leading to associated with localized FN signal loss beneath their cell body. Although matrix-specific cues are key drivers of tumor progression, how macrophages interact with and respond to matrix cues, and the molecular mechanisms underlying cell-matrix interactions remain poorly understood. Therefore, this platform provides a framework to identify matrix-specific therapeutic targets to modulate macrophage function and prevent tumor progression.

## Supporting information

Online Resource 1

## Acknowledgements

KW acknowledges startup funds from the Bioengineering Department at Temple University, an Office of the Vice President for Research Bridge grant, an Office of the Vice President for Research Blue Sky Initiative grant, and a grant (#2305) from the W.W. Smith Charitable Trust. GB acknowledges a Teaching Assistantship from the Bioengineering Department at Temple University. EB acknowledges a Diamond Research Scholarship. JL acknowledges a Teaching Assistantship. The authors thank the shared departmental resources at Temple University for equipment usage. The authors thank Kyle Shuler and Frederick Murphy from Nikon Instruments, Inc. for their assistance in acquiring images on their confocal.

## Statements and Declarations

The authors report no conflicts of interest.

## Authors’ contribution

GB: Conceptualization, Methodology, Validation, Data Analysis, Visualization, Writing. EB: Methodology, Validation, Data Analysis, Review, Editing. JL: Methodology, Validation. MSP: Methodology, Validation. KW: Conceptualization, Methodology, Validation, Visualization, Writing, Funding acquisition, Supervision.

