## Supplementary material for "Tumor-Associated EDA-FN-Enriched Matrix Instructs Macrophage Behavior": Online Resource 1

**Supplementary Figures:**

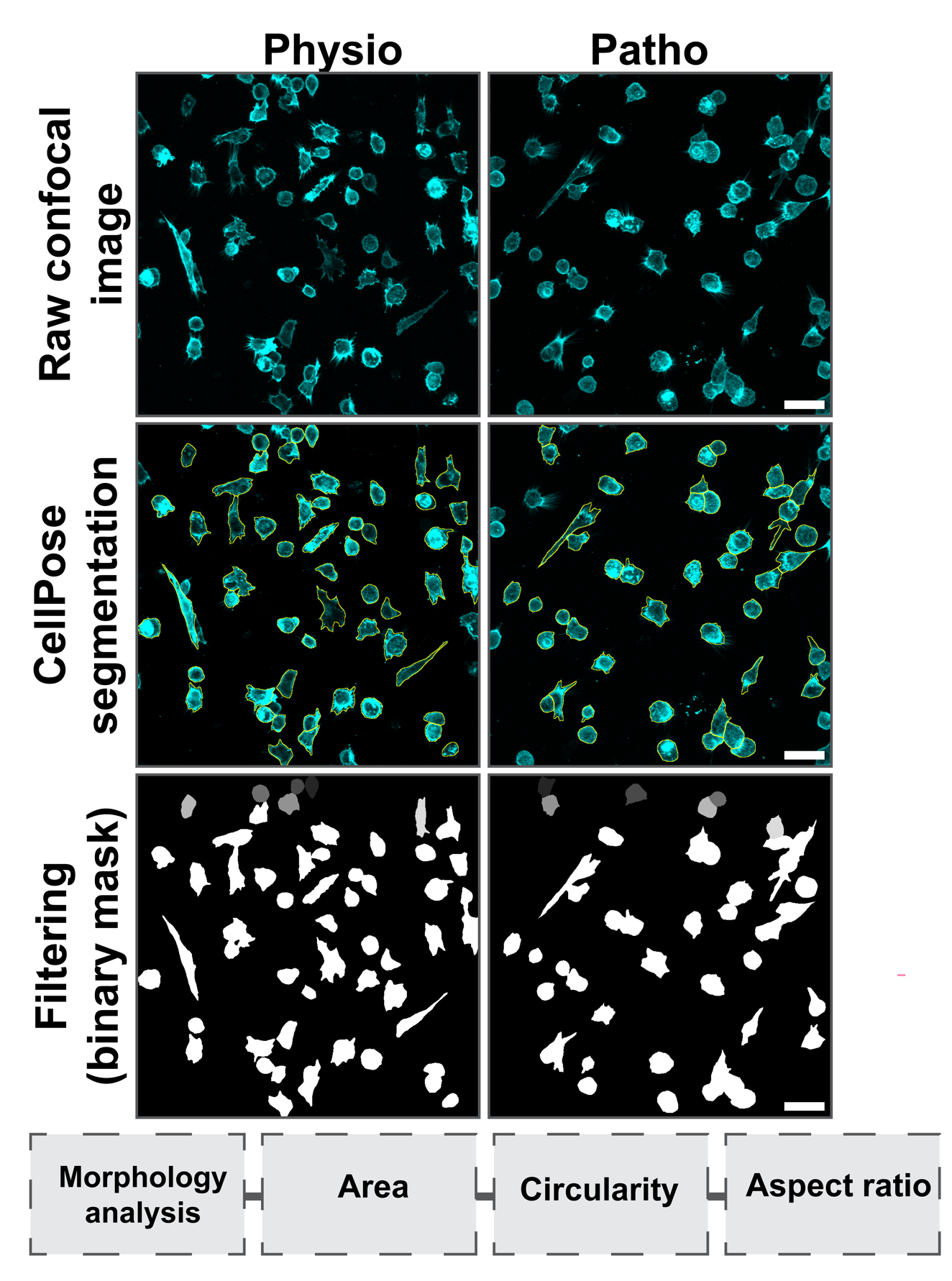

**Supplemental Fig. 1** Workflow for single-cell morphology analysis of macrophages. Representative confocal microscopy images of macrophages reseeded on physiological and pathological EDA-FN matrices. F-actin in cyan, CellPose single-cell segmentation shown as an outline overlay, and binary masks of macrophages following the removal of border-touching cells and cells with inaccurate segmentation masks. Scale bar, 50 µm.
